# Buzz-world: Global patterns and drivers of buzzing bees and poricidal plants

**DOI:** 10.1101/2024.03.06.583730

**Authors:** Avery L. Russell, Stephen L. Buchmann, John S. Ascher, Zhiheng Wang, Ricardo Kriebel, Diana D. Jolles, Michael C. Orr, Alice C. Hughes

## Abstract

Foraging behavior frequently plays a major role in driving the geographic distribution of animals. Buzzing to extract protein-rich pollen from flowers is a key foraging behavior used by bee species across at least 83 genera (these genera comprise ∼58% of all bee species). Although buzzing is widely recognized to affect the ecology and evolution of bees and flowering plants (e.g., buzz-pollinated flowers), global patterns and drivers of buzzing bee biogeography remain unexplored. Here, we investigate the global species distribution patterns within each bee family and how patterns and drivers differ with respect to buzzing bee species. We found that both distributional patterns and drivers of richness typically differed for buzzing species compared to hotspots for all bee species and when grouped by family. A major predictor of the distribution, but not species richness overall for buzzing members of four of the five major bee families included in analyses (Andrenidae, Halictidae, Colletidae and to a lesser extent, Apidae) was the richness of poricidal flowering plant species, which depend on buzzing bees for pollination. As poricidal plant richness was highest in areas with low wind and high aridity, we discuss how global hotspots of buzzing bee biodiversity are likely driven by both biogeographic factors and plant host availability. Whilst we explored global patterns with State-level data, higher resolution work is needed to explore local level drivers of patterns, but from a global perspective, buzz-pollinated plants clearly play a greater role in the ecology and evolution of buzzing bees than previously predicted.

## INTRODUCTION

Bees are one of the most important pollinator groups in both natural and agricultural systems (Winfree 2010; Orr et al. 2021). Yet in recent decades, there has been fervent discussion of potential bee declines and their causes (Cameron & Sadd 2020; LeCroy et al. 2020; Page & Williams 2023). Accordingly, understanding the dimensions of bee distribution is fundamental to targeting conservation efforts, and to developing strategies to respond effectively to anthropogenic threats including climate change, agricultural intensification, habitat loss, and urbanization (Potts et al. 2010; Cariveau & Winfree 2015). Furthermore, species vulnerability to environmental change is non-random and is linked to species-specific traits, yet for the majority of species the relationship between traits, habitat breadth, and distribution is only poorly understood.

Bees are very diverse, comprising over described 20,000 species spread among 508 genera and seven families (Ascher & Pickering 2023). However, like most invertebrates, bees generally lack the data required for high-resolution, global analysis (barring extensive modeling efforts; Orr et al. 2021, Dorey et al. 2023), and we have yet to explore how patterns of richness vary among families or how certain traits and their distribution over evolutionary time may influence these patterns. Current efforts to map large-scale patterns and drivers of bee distribution have focused strongly on the broadest phylogenetic patterns and abiotic drivers, but without substantial consideration of bee foraging behavior (Orr et al. 2021; Chesshire et al. 2022; Leclercq et al. 2022). This represents a fundamental gap in our understanding of the mechanisms underlying bee richness, because how and which flowering plant species bees interact with depends fundamentally on foraging behavior, which should in turn influence the geographic distributions of bees. One especially well-studied bee behavior that likely influences global patterns of bee distributions is the ability of many bees to vibrate flowers to extract their pollen (‘floral buzzing,’ ‘floral sonication,’ or ‘buzzing’), yet where and why this may be selected for, and even just how this varies geographically remains unknown. While buzzing is a widespread and potentially key driver of bee diversification (De Luca & Vallejo-Marin 2013; Cardinal et al. 2018), whether and how global patterns and drivers of bee distribution differ among bee taxa that can buzz versus cannot buzz is unknown.

Buzzing is performed by bee species from across all 7 bee families and at least 83 genera (these genera comprise ∼58% of all bee species; Cardinal et al. 2018; Table S1). Buzzing typically involves the bee biting the flower (usually the anthers), decoupling its flight muscles from its wings, and contracting these muscles rapidly to shake loose the pollen from the anthers (Buchmann 1983; Vallejo-Marin 2022; Vallejo-Marin & Russell 2023). While bees use buzzing to collect protein-rich pollen more effectively from many kinds of plant species (Buchmann 1985; Russell et al. 2017; Vallejo-Marin & Russell 2023), this behavior is most associated with extraction of pollen from so-called poricidal flowers – plant species that conceal pollen within tube-like morphology (typically the anthers). As a result, buzzing often enables access to these key floral resources when they would be otherwise inaccessible. Accordingly, buzzing is typically required for pollination of the more than 28,000 plant species across 87 families with poricidal flowers (an estimated 10% of flowering plant species; Buchmann 1983; Vallejo-Marin & Russell 2023; Russell et al. 2024). Poricidal plant species are also common in agriculture, and pollination of commercial crops such as tomatoes, cranberries, blueberries, and kiwis primarily depend on buzzing bees (Cooley & Vallejo-Marin 2021).

Given the importance of buzzing to bee and plant ecology and evolution, we hypothesize that this behavior should influence patterns of bee distributions and diversity, and may even be a significant driver of patterns in taxa where more species “buzz.” In particular, because bee taxa with a greater prevalence of buzzing can effectively access pollen from a broader range of hosts (Russell et al. 2017; Cardinal et al. 2018; Vallejo-Marin & Russell 2023), we predict that, globally, buzzing bee taxa should be more geographically widespread than taxa that do not buzz. Accordingly, assuming that all other key ecological drivers of buzzing and non-buzzing bee taxa are similar, regions with greater bee diversity should have a proportionally greater diversity of buzzing taxa. For instance, xeric and temperate zones are generally correlated with high bee species richness (Orr et al. 2021), and thus we would predict that bee taxa in these zones could therefore have a greater number of buzzing species than other zones, though this depends on the distribution of suitable hosts. Conversely, given that buzzing is often taxonomically restricted to bee clades with particular traits and habitats (e.g., buzzing is widespread among the mostly temperate bumble bees and rare among the sub/tropical stingless bees; Cardinal et al. 2018), the distribution and drivers of bee biodiversity may generally differ for buzzing and non-buzzing taxa.

In this study we map, model, and compare the known distribution of bee taxa reportedly capable of buzzing flowers vs total richness patterns. We examine the overall richness patterns of buzzing bees, and proportions of buzzing species within each bee family, and explore what the major drivers of these patterns may be, including the distribution of plant species with poricidal flower morphology to more clearly understand these patterns and why they may have developed.

## METHODS

This study involved the production of a global State-based inventory of bees, collation of key-traits of both bees (i.e., buzzing) and plants (i.e., if poricidal), the mapping of these traits, and finally an analysis of the drivers of both richness overall for each bee family, and the spatial drivers of buzzing.

### Bee species data preparation

Based on Orr et al. (2021), we used a higher-resolution version of the globally-authoritative DiscoverLife Bee World Checklist to link species to administrative areas (Ascher & Pickering 2023). Mapping was conducted at the State level (or regional equivalents) in most cases, and country level for some regions where sufficient State-based data did not exist; these designations are referred to as “administrative regions” hereafter. Species specific maps were developed for each species (totaling 20,941 species and 136,323 polygons representing the relationship between species and administrative regions) and could therefore be linked to traits for higher level analysis. Full methods for the compilation of species distribution are provided in Text S1.1 (see also supplementary materials in Orr et al. 2021). The all-bee analyses included all bee species from all families, but the family-level analyses excluded Stenotritidae and Melittidae, because these families are too small (21 and 213 species, respectively) for analyses to be meaningful, and because Stenotritidae is restricted to Australia.

### Coding buzzing

We grouped bees into four behavioral categories. Those in category 0 are assumed to never buzz flowers (no available published studies or expert observer evidence of buzzing) and 1-3 denote different scoring systems for buzzing bees (Table S1). In category 1, the strict approach, we considered only bee species known and published that buzz flowers to extract pollen. In category 2, or the consensus approach, we coded genera (and all species therein) as entirely buzzing only if they contained more than the median percent of buzzing species (9% across genera with bee species recorded as buzzing). In category 3, the liberal approach, we coded genera as entirely buzzing if they contained species documented to buzz flowers to extract pollen (this category was only used for a subset of analyses). Table S1 lists all bee species documented in the literature or observed by us or other bee researchers as buzzing flowers to extract pollen from flowers.

### Understanding patterns

Various drivers were considered to understand the overarching patterns of richness (Text S1.2; Table S2), the proportion of buzzing bee species and how this varies geographically. Firstly, we calculated the percentage of buzzing bees there were per family and for three regions (Americas, Europe-Africa, Asia-Australia). These values were averaged for each latitude with more than 10 species for each family and the three regions, then plotted.

We also ran cartograms for both overall richness and number of buzzing species. Cartograms basically calculate a global average richness per unit area, and then either grow or shrink each administrative area based on whether the chosen metric exceeds or falls below its global average (Figures S1-10). To quantify this in a standard way, we then calculated for each region what percentage increase or decrease the area showed and then both mapped the increases and decreases, and calculated the trends per group as averages.

### Drivers of buzzing

We considered 52 different variables as potential drivers of taxonomic diversity and buzzing (see Table S2), and for each of these included maximum, minimum, mean, and standard deviation within each administrative area, with the exception of plant richness (just maximum and mean) and poricidal plant richness (total per administrative area). These variables included different elements of climate, as climate and its facets are likely primary drivers of buzzing, including possibly via their impacts on plants. Environmental variables were selected based on their potential to drive richness patterns, and our previous analysis (Orr et al. 2021).

Poricidal plant species were mapped using the species-specific plant data compiled by Liu et al (2023), with some additional species added from the Kew Plants of the World Database (accessed October 2023). The list of poricidal plant genera is provided in Table S3, and all genera were treated as monomorphic for poricidal state following Russell et al. (2024). For a complete description of how poricidal plants were characterized, see supplementary material and Russell et al. (2024). Full details on the compilation of poricidal plant maps is provided in Text S1.2. In addition, we used averages of plant richness based on Denelle et al. (2023). These factors were then evaluated for their role in driving all bee – and buzzing bee – distributional patterns, as detailed below.

### Analyzing trends

General linear models were run in Spatial Analysis for Macroecology (SAM), all based on the same administrative areas as above. First, we ran single regressions for overall richness (as well as percent buzzing) with each variable, then we used this to make the best model by using initially the variables with the strongest relationships with richness for the group, and sequentially removing variables which were not significant in the model until all variables were significant. We did not analyze the drivers for buzzing overall for all bees, given that drivers may vary by family; thus including them all within a single analysis would be weighted by the most species-rich families. We then ran the regression analysis within ArcMap (ArcGIS, Esri) using the spatial statistic tools, and regression analysis and an exploratory regression of the preselected variables, as well as exploring the drivers of poricidal plant richness.

Three versions of models were run for each family. The first used the variables selected for overall species richness within each group, the second was for buzzing species (using the percentage of species coded as 1 or 2 for buzzing), and the third was for total richness patterns, but using the variables for buzzing richness, to understand if the drivers for buzzing differ from that of overall richness. We assessed both the main, and most explanatory variables for each of the three scenarios for each family, then mapped the drivers of poricidal angiosperm richness. We also explored if average flower size was smaller for poricidal plant species using the same list of poricidal genera and the dataset on plant traits from Song et al (2022) (Table S4).

## RESULTS

### Patterns of bee and trait richness

Patterns of overall bee species richness were largely consistent with previous analyses (Orr et al. 2021), with richness peaking in North America (especially in the Southwestern United States), and also areas of the Middle East, Southern South Africa, and Australia (Figure 1). However, the proportions of bee species from each family varied by region. For example, in northern North American states, over 50% of bee species were halictids (Figure S1), but with decreasing latitude, the ratio of Apidae to Halictidae increases so that Apidae eventually becomes the dominant group in terms of species richness. This pattern is similar in South America, with Halictidae dominating in the southeast, followed by Colletidae. In Africa-Europe (Figure S4), Halictidae overall dominates, followed by Apidae, then Colletidae, but Halictidae dominates in equatorial regions in part given Nomiinae bee dominance, whereas Apidae becomes dominant in southeast Africa. In Asia, Apidae dominates in northern latitudes, but Halictidae becomes more dominant at lower latitudes, in addition to Australia (Colletidae becomes more diverse in Australia-New Zealand) (Figure S7). Colletidae also generally holds a larger share in the Southern Hemisphere, especially Australia. Thus, global patterns should generally be broken down by region and family, as patterns (and likely their drivers) are clearly different between families and regions.

**Figure 1.**
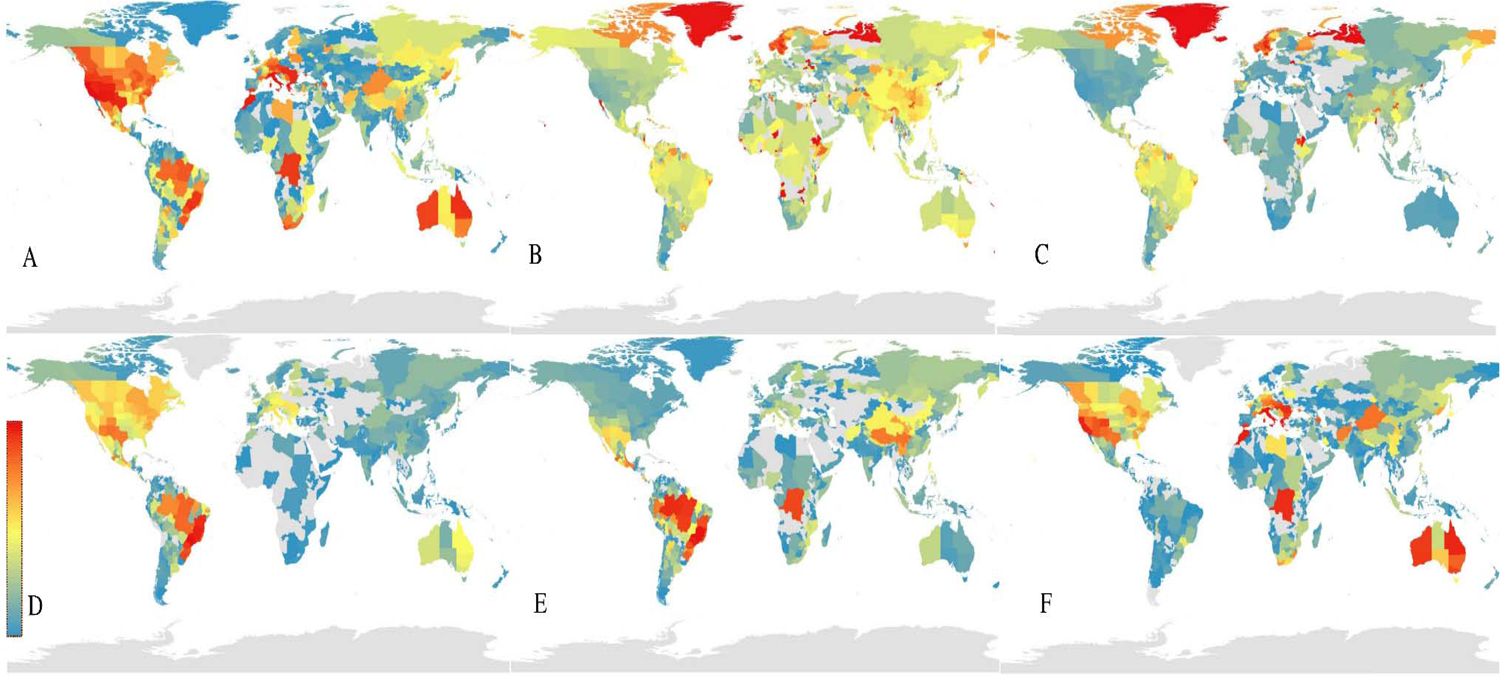
Patterns of richness across regions for all bees and for buzzing bees. Redder colors indicate higher values, blue colors indicate lower values, and grey indicates no species recorded. A. Total bee species richness (maximum richness value 1619). B. Percentage of bee species in each region that buzz (categories 1+2+3, the most liberal coding). C. Percentage of bee species that buzz (categories 1+2). D. Number of buzzing bee species, coded category 1 (maximum 108). E. Number of buzzing bee species across categories 1+2 (maximum 238). F. Number of buzzing bee species, across categories 1+2+3 (maximum 331).

Patterns for buzzing species depart from the overall patterns of richness. In Andrenidae, the patterns in North America (Figure S2-3) and Europe are similar (Figure S5-6), however a greater percentage of species buzz in South America where there are many buzzing panurgine bees. In contrast, a much smaller percentage buzz in East Asia or North Africa (where non-buzzing *Andrena* dominates), though these show latitudinal gradients which differ by region. Conversely, in Apidae, patterns are overall similar in most regions, but a greater percentage of species buzz in Northeast Asia (Figure 2), and fewer species buzz in parts of Southern and Southeast Asia (Figure S8-9). For Colletidae, patterns are quite similar between buzzing and non-buzzing species. Halictidae also show similar patterns, though fewer species buzz in Western South America. Megachilidae show some distinct differences, with very few species in the Southern Hemisphere buzzing and more temperate species buzzing.

**Figure 2.**
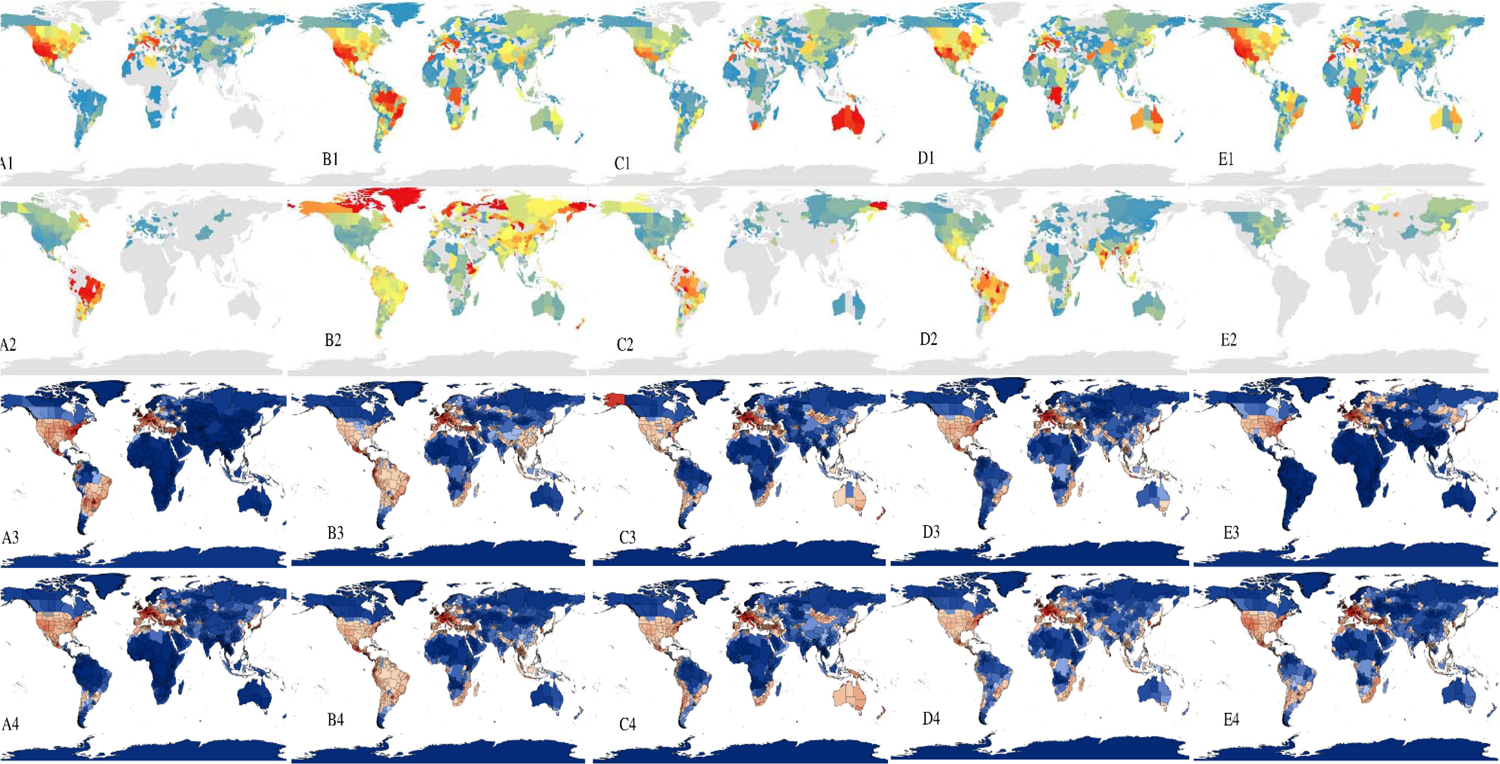
Patterns of richness by family, with redder colors indicating higher values, bluer colors indicating lower values, and gray color indicates no recorded. A. Andrenidae (max richness 548), B. Apidae (max 388), C. Colletidae (max 334), D. Halictidae (max 212), E. Megachilidae (max 400). Numerical ordering is 1) richness of each family, 2) percentage of buzzing species (categories 1+2), 3) areas with more species than expected per unit area relative to the global average, and 4) areas with more or less species per unit area relative to the global average.

When the percentage of species buzzing per family and region are plotted, different patterns emerge, and these vary depending on how stringent the definition of buzzing is. In the Americas, Apidae and Halictidae buzzing peaks in tropical regions (Figure S3), where up to around 60% of species buzz in tropical regions using the stricter definition of buzzing (categories 1+2), decreasing at higher latitudes (Figure 2; Figure S2). Colletidae also shows this pattern, but not as strongly (and with lower diversity in the tropics), while Andrenidae exhibits greater proportions of buzzing at higher latitudes. These patterns shift if a liberal definition of buzzing is used (categories 1+2+3), such that while Apidae still shows a tropical peak in the proportion of bees that buzz (Figure S3), Halictidae tends to increase in the Northern Hemisphere, where richness is also higher, but patterns in other groups change less; thus, a stricter definition was used when trying to understand drivers of the buzzing trait.

In the Europe-Africa region, Halictidae shows a slight peak in the proportion of bees that buzz in equatorial regions (categories 1+2), but Apidae peaks in buzzing in northern latitudes in Europe (Figure 2; Figure S5, S8), and Andrenidae and Colletidae peak slightly in the north, though largely due to low diversity elsewhere. These patterns somewhat disappear if the liberal categorization (1+2+3) is used, though Colletidae shows a much higher percent increase in buzzing in northern latitudes (Figure S6).

In the Asia-Australia region, Apidae shows the same pattern as in the Europe-Africa region (with increasing percentages of buzzing in northern latitudes due likely to increasing relative richness of *Bombus* and perhaps *Lasioglossum* species, both when stricter buzzing categories (1+2) (Figure S8) and the most liberal buzzing categorization (1+2+3) are considered (Figure S9). Halictidae increases in the percentage of buzzing in equatorial regions when using the stricter categories and there is no strong pattern when the liberal buzz categorization (1+2+3) is considered. Conversely, the percentage of buzzing Colletidae species increases in higher latitudes in both the Northern and Southern Hemispheres when the liberal categorization is considered (but only very minor increases in these regions when stricter buzz categories are considered). Megachilidae shows a slight increase in percent buzzing in high latitudes in the Northern Hemisphere (Figure 2).

Some of these regional differences in patterns may reflect the distribution of particular genera with high proportions of buzzing. For example, *Bombus* is a well-studied and widely distributed group of buzzing species, with many of its species in the neotropical regions and few in the paleotropics (Figure S10). This trend, and the regional and latitudinal patterns in *Bombus* may be partially responsible for the patterns in percentage of buzzing Apidae species across higher latitudes of Europe, where they are more prevalent and make up a bigger proportion of bee species, and also for Asia (especially the Himalayan region, where *Bombus* are particularly speciose; Figure S10).

### Drivers

There were generally distinct and important differences between models for the percentage of buzzing species and bee overall richness for each family (all regression results are provided in Table S5, see also Text S2.1 for a detailed breakdown of drivers by family). For three of the five families with sufficient diversity and spatial spread for global analysis (Andrenidae, Colletidae, and Halictidae), the richness of poricidal plants was the main variable driving higher percentages of buzzing species, despite the fact that it was only a very weak driver for total richness in those families (Table S5), demonstrating a link between buzzing species biodiversity and the plants these bees use. The highly diverse family Apidae also exhibited a positive relationship between poricidal plant richness and buzzing species richness, but there was no relationship of poricidal richness with overall Apidae richness. For Megachilidae, there were no significant relationships with poricidal plant richness, however we recognize that only a small fraction of species in this family have been observed to buzz flowers (8 of 4169 total species).

The drivers differed by bee family, and for overall richness versus the percentage of buzzing species (Table S5; Text S2.1). These differences may also be amplified in different regions based on species biogeography and prevailing conditions (Almeida et al. 2023). Given the high importance of poricidal richness for buzzing in many bee taxa, we then explored if poricidal richness might be driven by elements of arid climates, given its importance to bee richness (Orr et al. 2021). Unsurprisingly, total plant species richness was a major correlate of poricidal plant species richness, explaining why poricidal richness rarely appeared in most overall bee species richness models, but was in most buzzing species richness models. The next most important correlate for poricidal plant species richness was a negative relationship with continentality and then a strong negative relationship with latitude. Poricidal richness was overall associated with higher plant richness, lower latitudes, low continentality, low wind, high aridity and high evapotranspiration. The value of poricidal richness in many models of buzz percentage is likely to have reduced model reliance on co-occurring factors of higher aridity and low wind that co-occur with higher poricidal richness. Furthermore, in terms of mean values, the average altitude and sampling aridity of poricidal vs non-poricidal plants was very similar (1398.5m-1393.6m; 4.07-4.3), while poricidal plants had a slightly higher mean latitude (18.5° for non-poricidal vs 20.9° for poricidal), and the mean flower-size was slightly smaller in poricidal species (3.15 vs 2.98 cm).

## DISCUSSION

In this study, we explore the environmental conditions that have facilitated the evolution and biogeography of buzzing bees and poricidal flowers. Patterns of overall bee diversity differ from patterns of richness for buzzing bees, with patterns varying across regions among bee families and differences particularly pronounced in some bee families. In the Americas, overall richness peaks in North America, but the percentage of species that buzz is disproportionately high in temperate regions (latitudes 40 and −40) for Andrenidae, Colletidae (latitudes 20/-20-40/-40), and Megachilidae (latitudes 30-60), whereas Apidae and Halictidae largely show peak proportion buzzing in equatorial regions, though Apidae also shows a peak at higher latitudes in part due to *Bombus* (Figures S1-10). Patterns in Europe-Africa echo those of the Americas, although the peak in Europe for proportion of species that buzz is even more pronounced (Figure S1-9). Asia-Australia shows a somewhat different pattern, with stronger peaks of proportion buzzing in temperate regions (for all groups except Apidae), but with proportion buzzing peaking even more strongly in temperate Northern regions. In most families, the drivers of richness for buzzing species were different from those of overall bee richness, and in many cases if the model was optimized for buzzing species there was not a significant relationship with overall plant species richness. This highlights that buzzing has been selected for and persists in certain conditions, but that these conditions vary between different regions and taxa. Overall, we find that poricidal richness is one of the main predictors for the global biogeography of buzzing bee richness for most bee families.

Our results indicate that poricidal plant richness plays a key role in driving the percentage of bee species that buzz flowers for the Andrenidae, Halictidae, and Colletidae, and to a lesser, but still significant extent, the Apidae. This is surprising, because most buzzing bees are considered to be relatively generalized pollen foragers using a wider variety of plants compared to floral specialists that focus on typically few related species (e.g., Houston & Ladd 2002; Schlindwein 2004; Corbet et al. 2014; González-Vanegas et al. 2021) and are often observed collecting pollen from both poricidal and non-poricidal floral resources (e.g., Mesquita-Neto et al. 2018; Houston & Ladd 2002). In addition, although buzzing bees dominate access to pollen from poricidal flowers (Vallejo-Marin & Russell 2023), buzzing behavior is not specialized to poricidal floral morphology and is in fact used by bees to extract pollen from poricidal and non-poricidal plant species (Buchmann 1985; Russell et al. 2017). One simple explanation for this discrepancy is that buzzing bees may be more closely associated with poricidal flowers than presently understood, or that dominance of poricidal plants in some regions may exclude bee species that do not buzz. Much of the research on foraging associations between buzzing bees and flowering plants focuses on temperate regions, but buzzing bees in tropical regions often forage substantially on poricidal hosts (e.g., Mesquita-Neto et al. 2018; Vit et al. 2018; González-Vanegas et al. 2021; Delgado et al. 2023; Pemberton 2023). Given that poricidal species richness is highest in the tropics and plant species richness is often strongly positively associated with plant abundance (e.g., Bock et al. 2007; Sutter et al. 2017; Oloffson & Shams 2007), poricidal flowers are likely an abundant resource for tropical buzzing bees such as some sweat bees (Augochlorini) and orchid bees (Euglossini) (Mesquita-Neto et al. 2018; Delgado et al. 2023; Pemberton 2023). Our results also show this, with high poricidal richness and buzzing bee richness in some tropical regions, such as the Amazon (Figures 1, 2). More generally, poricidal plants can offer particularly protein-rich pollen (Roulston & Cane 2000) and produce substantially more pollen per flower than non-poricidal species (Buchamann 1983; Russell & Papaj 2016). Additionally, while pollen is the principal food reward offered by poricidal flowers to bees in exchange for pollination, some non-poricidal species may deter bees from actively collecting pollen via chemical defenses (Irwin et al. 2014; Vanderplanck et al 2018; Palmer-Young et al. 2018). Nonetheless, little is known about the nutritional or metabolic advantages to buzzing bees of foraging on poricidal flowers or the relative importance of poricidal hosts for most buzzing bee species.

Both hotspots and drivers of bee richness varied among families, with different latitudinal patterns among regions likely resulting from both biogeographic factors and niche availability. Furthermore, for highly diverse bee families, drivers likely differ among genera, which should drive distributional differences among regions. Additionally, buzzing behavior is rarely reported in particular groups, such as in the Andrenidae (20 buzzing species documented across 14 genera; out of 3089 total species across 69 genera) and the Megachilidae (8 buzzing species documented across 3 genera; out of 4169 total species across 95 genera) (DiscoverLife; Cardinal et al. 2018). Accordingly, it is possible that underreporting of buzzing species could thus explain the lack of an association between buzzing megachilids and poricidal richness. On the other hand, buzzing species may truly be a rare occurrence in the Megachilidae, for instance. In contrast, buzzing behavior is well documented in the largest family Apidae, and the reduced importance of poricidal flowers in buzzing Apidae biogeography is likely legitimate and may be due to the diverse life-histories within the group. For instance, socially-parasitic species (kleptoparasites), which do not collect pollen from flowers, comprise nearly a quarter of the Apidae (Cardinal et al. 2010). Eusocial species that are extreme generalists and forage from dozens or hundreds of non-poricidal (and poricidal) plant species are prevalent in Apidae and might also dilute any overall association between buzzing and poricidal richness, which warrants further exploration.

The evolution of poricidal flower morphology is frequently attributed to selection to reduce pollen loss by less efficient generalized pollinators, as well as to selection by buzzing bees (Buchmann 1983; De Luca & Vallejo-Marin 2013; van der Kooi et al. 2021; Vallejo-Marin & Russell 2023). Our results suggest that poricidal morphology is an adaptation to certain climates and abiotic conditions. Notably, poricidal richness was positively associated with high aridity and evapotranspiration, suggesting that poricidal flowers, which enclose pollen within tube-like morphology, might be an adaptation associated with keeping pollen viable in desiccating environments (Vallejo-Marin & Russell 2023; Russell et al. 2024).

Our results also indicate that how environment and pollinator behavior interact is likely a key factor in the evolution of poricidal plants. During buzz pollination, pollen is ejected from poricidal morphology into the air, before landing on the bee (Buchmann & Hurley 1978; Corbet et al. 1982; Buchmann 1983). Windy conditions might therefore reduce the likelihood of pollen transfer to the bee, thus reducing the selective advantage of buzz pollination relative to other pollination systems. Consistent with this hypothesis, we found that poricidal richness was negatively associated with high wind. Similarly, the positive association between increasing aridity and poricidal plant richness may be in part due to associated negative effects of aridity on flower size (Galen et al. 1987; Galen 1999; Xie et al. 2016; Lozada-Gobilard et al. 2022). Pollinator specialization, which tends to enhance pollination success and drives angiosperm evolution (Armbruster 2014, 2016; Brosi 2016; Villalobos et al. 2019), also often increases with decreasing poricidal flower size (Delgado et al. 2023. Although how behavior and environment interact to drive poricidal flower evolution has not been studied to our knowledge, such interactions are known in other pollinations mutualisms. For instance, selection for ultraviolet (UV) absorbing patterns of high-altitude flowers is likely often driven by both higher UV irradiance and UV preferences of pollinators (e.g., Koski & Ashman 2013, 2015; Klomberg et al. 2019; Koski et al. 2020).

In conclusion, the percentage of bee species that buzz flowers is clearly frequently driven by poricidal plant species patterns (see also Pacheco Filho et al. 2015), which are in turn driven by aridity and wind. Accordingly, these traits are interlinked and reflect broad-scale environmental patterns. Furthermore, both hotspots and drivers of richness varied for different bee families, but both spatially and taxonomically higher resolution work is needed to explore local level drivers of these patterns, particularly in rapidly changing environments such as mountain landscapes (e.g., Knight 2022). A major gap in our understanding of these patterns remains the underreporting of buzzing bee species, particularly in less sampled and remote geographic regions (Cardinal et al. 2018; Orr et al. 2021). This weakness is especially evident when examining hotspots of poricidal plant species richness: many tropical regions with high poricidal richness have a lower richness of recorded buzzing species. Further work will be required to determine whether these patterns are also the result of regional differences in the ecological drivers of buzzing. Additionally, while our focus here lies in patterns of species richness, patterns of species abundance have obvious importance to understanding biogeographical patterns and should therefore be a major focus of future work examining the (co)occurrence of buzzing bees and poricidal plants. Altogether, the results here represent a major step towards understanding the ecological and evolutionary scenarios in which buzz pollination evolves and persists.

## AUTHOR CONTRIBUTIONS

ACH, ALR, SLB, and MCO conceived the study. ACH, JSA, and MCO collected the bee data. ALR and RK provided the list of poricidal species, DDJ helped put together the initial list of poricidal species, ALR, MCO, and JSA listed the buzzing traits. ACH mapped the distributions, developed the methods and analyzed the data. ZHW provided the plant distribution data. ACH, ALR and MCO wrote the original draft of the manuscript. All authors revised the paper and approved submission.

## ACKNOWLEDGEMENTS

We acknowledge that work performed by ALR occurred on unceded traditional territory of the Kiikaapoi, Sioux, and Osage. MCO acknowledges the President’s International Fellowship Initiative Visiting Scientist program (2024PVC0046).

## CONSENT TO PARTICIPATE

Not applicable

## CONSENT FOR PUBLICATION

Not applicable

## DATA ACCESSIBILITY

Datasets supporting this article are available as electronic supplementary material and bee richness distributions are available via DiscoverLife and the supplemental materials of Orr et al. (2021).

## SUPPLEMENTAL MATERIAL

### SUPPLEMENTAL TEXT

#### Text S1: Supplemental methods

##### S1.1. Mapping bee distributions

Compared to prior efforts to map bees, we increased resolution and used State/province or major island level for most areas. We used the Global Administrative Areas database as a reference (GADM). However, in this study we directly linked species to administrative areas, creating a much larger file, so every species could be separately mapped. We iteratively converted all state-based notations into HASC codes to ensure the country codes were all coded as ISO2 and not FIPS numbers. First, we used the entire checklist and summarized it to provide a list of state and country names, so we had a much easier reference for conversion. All codes and names from the original checklist file were converted to HASC codes, by first downloading all HASC codes with state-names (from Statoids) and then using join and relate in ArcMap 10.8 to connect the inventory of areas noted for species. We also noted if the region referred to was in multiple different regions, and manually corrected such HASC codes when needed (e.g., there are multiple San Jose’s). If multiple names fell within one HASC region, they were given the same code, and then duplicates of the same species to the same state were removed. Statoids was used as a reference for converting HASCs. An exception to this was in areas with big states/provinces and good data, such as France, where species were often listed at finer scales, and countries like the United Kingdom where the listing system is slightly different within HASC and counties are exceedingly small. Conversely, some areas had very under-collected regions. Thus, for biogeographically complex regions that include a combination of islands and provinces, as well as for regions that were listed in a range of ways (Indonesia and the Philippines), new island-based divisions were created and species lists corrected as appropriate. Furthermore, for some regions (such as various African countries) inventories are known to be incomplete, thus such countries were left at the country level and a separate list created for state vs country-based listings. Two separate shapefiles were also created to separate countries with enough data for state-level analysis and those that needed to be examined on a country-level.

Once all codes, names and notations were converted to HASC codes, they were cross-linked to the species checklist and then connected to the GADM layer. This gave a total of 136,323 links between the state-based administrative areas and species distributions.

A list of bee species that buzz was created separately (see next section for details), and we assessed the number of species with the three categories of “buzz” state, and the percentages for different categories and overall, for all regions. This included 14,672 non-buzzing species, 410 species included as “1” for buzzing (the strictest buzzing category), 2, 1674 species included in category 2, and 4185 species included in category 3. In addition, we separated these analyses by bee family so we could differentiate patterns that may reflect the different evolutionary and biogeographic histories of different groups, as the percentage of buzzing species does vary by group.

##### Text S1.2 Mapping plant and poricidal plant richness

Plant richness was calculated by taking the mean and maximum richness of the Ensemble_Prediction_7774_Eckert-IV from the GIFT shiny app (https://gift.uni-goettingen.de/shiny/predictions/). This was then integrated with a stencil of our administrative regions using the Intersect tool in ArcMap to map both the maximum and mean richness per administrative area. For poricidal plant richness, we first created a list of poricidal genera (Table S3) from Russell et al. (2024) and then extracted the species ranges for all species referenced in the Peking University database (Liu et al. 2023; https://en.geodata.pku.edu.cn/index.php?c=content&a=list&catid=199). The list was generated at the genus level, because numerous well characterized genera are frequently monomorphic for the presence/absence of poricidal morphology (e.g., *Solanum*, *Senna*, *Miconia*) and information for all species in rarer and/or tropical genera is frequently sparse. However, we recognize that monomorphism within genera should not be taken for granted and there is evidence of potential exceptions (e.g., Renner, 1989; Gavrutenko et al., 2020). We also confirmed non-poricidal morphology for monomorphic genera when morphological information was available. See Russell et al. (2024) for a complete description of the methods used to find and characterize poricidal plants. Five additional genera were later appended to the list, with their species extracted from the Kew Plants of the World Online (https://powo.science.kew.org/). We used the PKU database as not only is it carefully checked, but the completeness and certainty of data has been mapped (which is not the case for other global checklists) and efforts have been made to increase completeness in regions like the African continent, where there are major data-gaps in the GIFT checklist. As the regions did not exactly align between the Kew Database and those used by the PKU team, administrative units were intersected in ArcMap with the stencil used by the PKU team and the Kew maps to ensure all species were mapped accurately.

Unfortunately, the number of administrative units varied between our administrative map and that used by the PKU group. Thus, having mapped the richness of poricidal plants for the PKU stencil (using the summary statistics tool in ArcMap), we then used a spatial join where the PKU stencil units were larger than our existing stencil. For areas where several units within the PKU units fitted within our administrative units (i.e., parts of the Amazon, much of Australia), we extracted the species maps for these regions, intersected them with our administrative stencil, then used the summary statistics to first list and then count the species within each of our administrative units.

#### TEXT S2: SUPPLEMENTAL RESULTS

##### Text 2.1 Drivers of richness

In Andrenidae, we see major differences in the drivers between buzzing and overall richness. The top variable, which showed a significant positive relationship in 100% of instances, was the richness of poricidal species (Figure 3), but this was only the case for buzzing species (percentage of species classed as 1-2), conversely the top variable for total richness with buzz variables was latitude, and for total richness with optimized buzz variables was a negative relationship with count of months over 10°C. From all analyses, the most important variable for Andrenidae for buzzing species was poricidal richness; this was followed by latitude, which had a significant negative relationship in all models. Conversely, overall andrenid richness had a strong positive relationship with latitude, indicating that in areas with fewer species (at lower latitudes), more species buzz. For buzzing species, the number of months with mean temp greater than 10°C also had a positive relationship with buzzing in all cases, but a negative relationship with total richness. Continentality and mean evapotranspiration also showed a negative relationship with buzzing percentage (Table S5), but only weak relationships with total richness (and the relationship with continentality was positive). The most important driver for richness overall was latitude (positive) followed by a strong positive relationship with precipitation of driest quarter (which was also positive for percentage buzzing). Overall, while more andrenid species occur in higher (temperate) latitudes with higher moisture during the driest parts of the year, buzzing peaked at lower latitudes.

**Figure 3.**
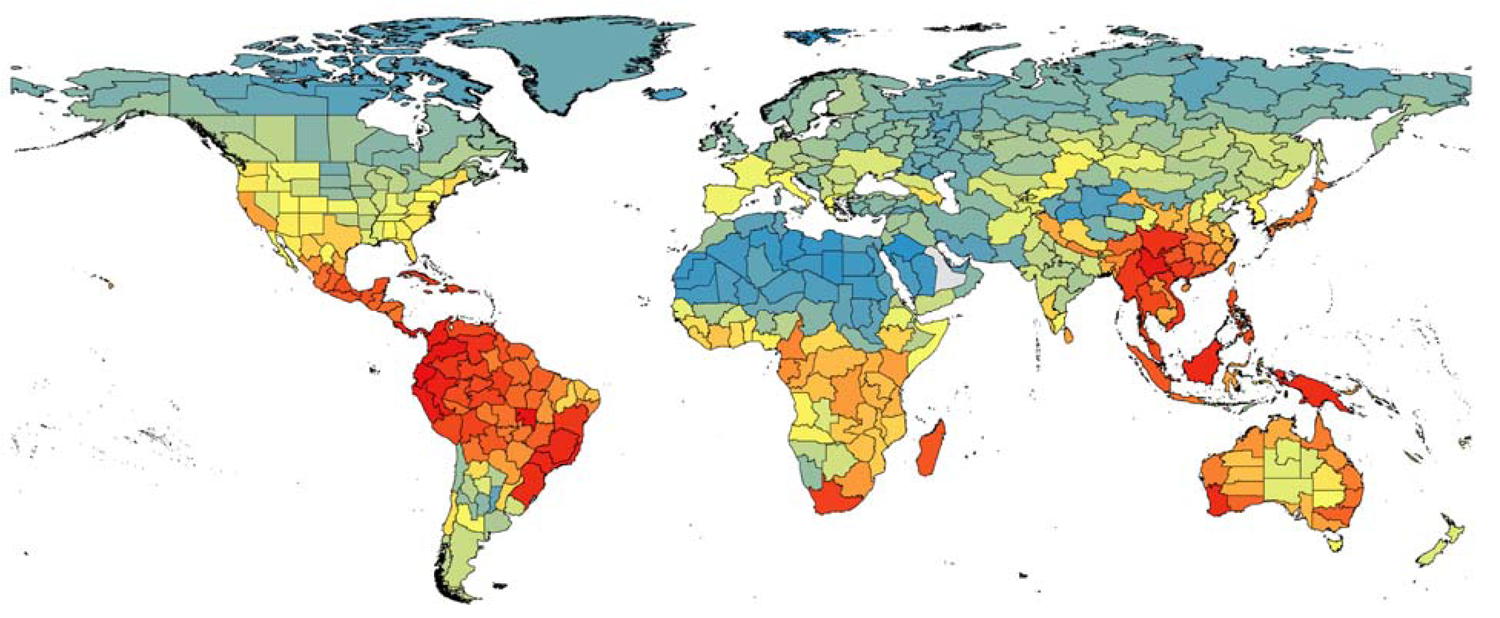
Patterns of richness in poricidal plant species across regions, maximum richness 2484.

For Apidae, while buzzing was less impacted by poricidal species richness, there was a strong negative relationship with maximum annual temperature (whereas total richness had a weak relationship with maximum temperature). Climate moisture index also had a consistently positive relationship with the percentage of buzzing species, as well as richness overall. Mean temperature of the coldest quarter had a negative relationship with the percentage of buzzing species, so areas with higher temperatures during the coldest parts of the year had smaller proportions of buzzing species, but this was more ambiguous for total richness. Poricidal richness showed the next (4th) most consistent (and positive) relationship with percentage of buzzing species, as well as for richness overall. Areas with high temperature seasonality also had a higher percentage of buzzing species and richness overall. Buzzing percentage also decreased with higher latitudes, but did not have a strong relationship with richness overall. Overall richness had an interesting relationship with aridity, showing a negative relationship with the Thornwaite aridity index, but a positive relationship with other metrics of maximum aridity; demonstrating a nuanced, but important impact from aridity.

Colletidae also showed poricidal richness as having the most consistent, and positive relationships with percentage buzzing, but only a weak contribution to overall richness (Table S5). The number of months with mean temp greater than 10L also showed a consistent positive relationship with buzzing, but a negative relationship with richness overall. Potential evapotranspiration seasonality had a negative relationship with percentage of buzzing, but a strongly positive relationship with richness overall. Mean monthly potential evapotranspiration (PET) of the wettest quarter had a consistent positive relationship with the percentage buzzing as well as richness overall. Plant richness overall was the next most important variable for percentage buzzing in Colletidae, and has a positive relationship, though this is more variable than with richness. For overall colletid richness, PET seasonality has a strong negative relationship, whereas there was no consistent relationship with the percentage buzzing. Maximum solar radiation has a strong positive relationship with richness. Mean temperature of the wettest quarter also had a strong and consistent relationship with richness, which was followed by mean monthly PET of the warmest quarter (also with a positive relationship).

For Halictidae, poricidal richness had a strongly positive relationship with buzzing percentage, but none with overall richness. This was followed by mean temperature of the wettest quarter which had a positive relationship with buzzing, but no relationship with richness overall. Percentage of buzzing species showed a negative relationship with annual temperature range, but a slight positive relationship with richness overall. Minimum temperature of the warmest month also had a negative relationship with the percentage of buzzing species, and a positive relationship with richness overall. Mean evapotranspiration also showed a negative relationship with percentage of buzzing species, but a positive relationship with richness. For overall richness minimum isothermality was the most consistent variable, but showed a negative relationship with richness. This was followed by maximum aridity index, which showed a positive relationship with richness, but no relationship with buzzing. Maximum mean diurnal range has a positive relationship with richness, but no relationship with buzzing. Plant richness had the next most consistent positive relationship with halictid richness.

For Megachilidae, mean wind had the most consistent relationship and positive relationship with the percentage buzzing, as well as a weaker, but positive relationship with richness overall. Latitude shows a positive relationship with buzzing, but a less consistent relationship with richness overall. Minimum precipitation of driest month also had a positive relationship with the percentage buzzing, but no relationship with richness overall. Growing degree days 5 shows a strong negative relationship with the percentage of buzzing species, but no relationship with richness overall. Maximum evapotranspiration also has a negative relationship with buzzing percentage, but a positive relationship with richness. For Megachilidae richness overall mean diurnal temperature range has a strong positive relationship, as did potential evaporation seasonality. Minimum solar-radiation shows a strong negative relationship with richness, but a positive relationship with mean temperature of wettest quarter and plant richness.

###### Latitudinal patterns

For plots of species richness, where percentage of species that buzz is calculated, areas with under 10 species were excluded, as percentages in values below this are likely to be highly stochastic. For example, 100% of 1 is 1, so if only a single species is present and did or did not buzz, comparing that 100% or 0% to more species-rich areas would not be meaningful or representative. Averages were taken for all regions at each latitude for each of the three global areas to examine overall patterns.

###### Americas

**Figure S1:**
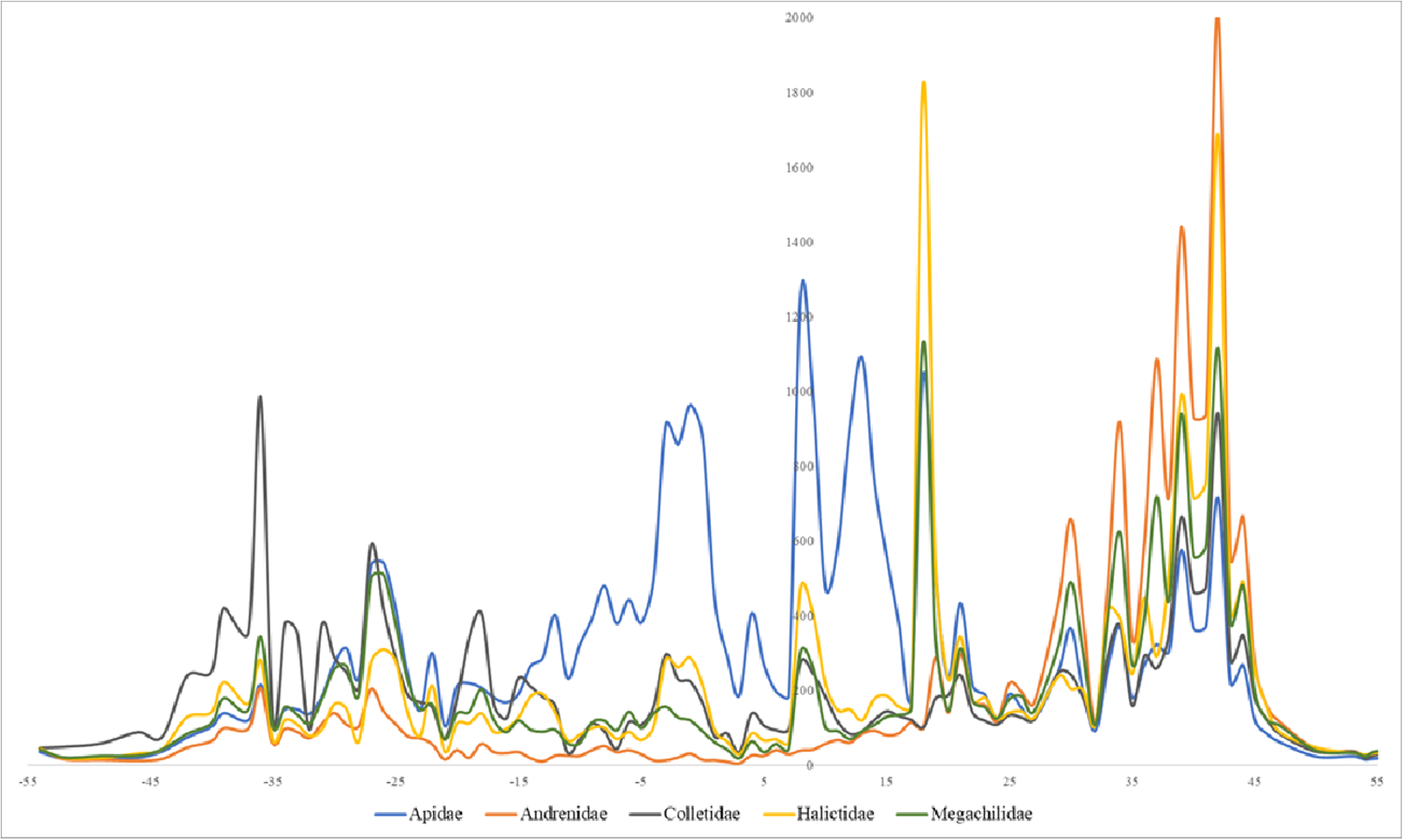
The degree in distortion of cartograms accounting for richness overall in the Americas, highlighting areas with more than the global average richness per unit area per family. The temperate zone peak indicates disproportionally high diversity for multiple families, whereas Apidae has more of a peak in more equatorial regions.

**Figure S2:**
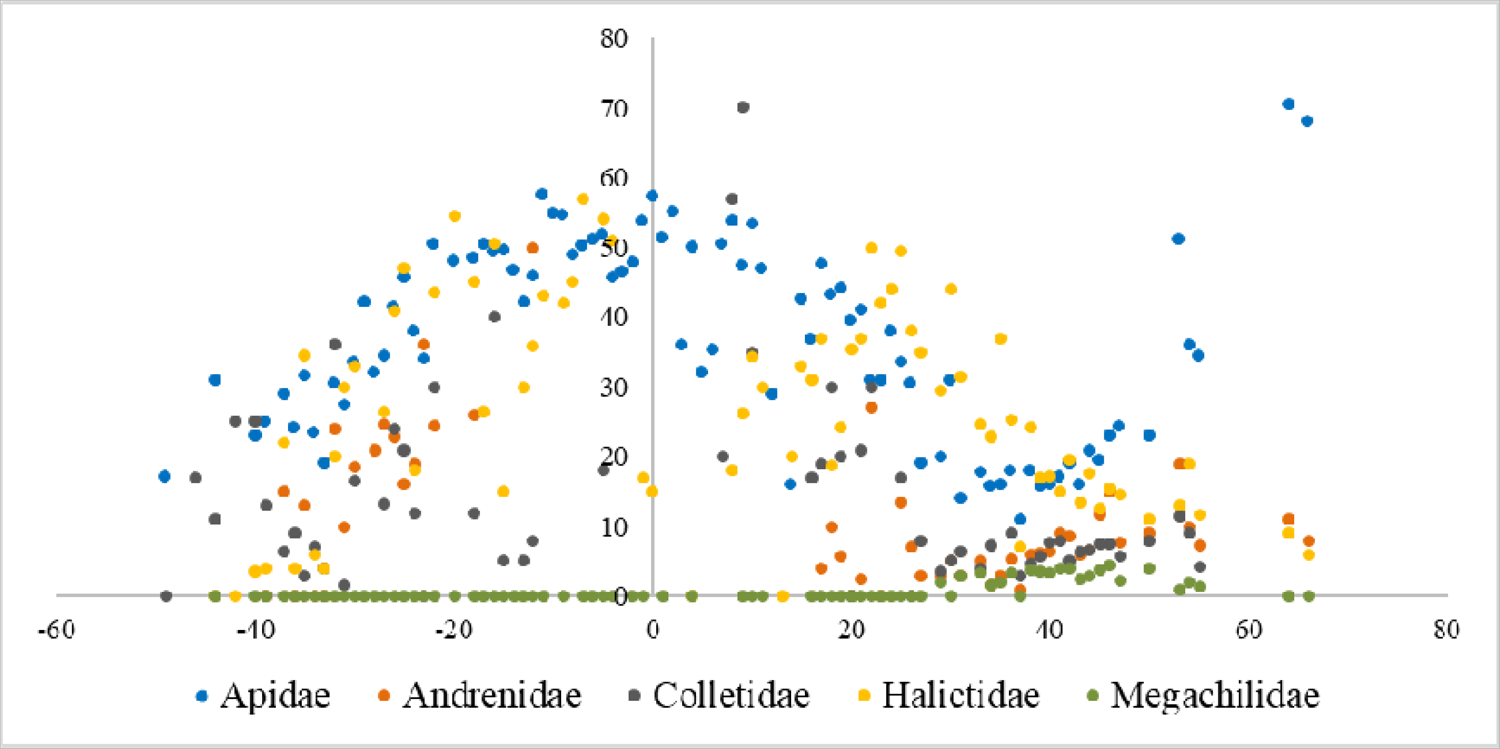
Percentage of species that buzz (categories 1+2) on average with latitude, including all areas with more than 10 species in the Americas. Very strong tropical peaking in the percentage of buzzing in Apidae and Halictidae. Whereas for Colletidae, peak percentage is higher in temperate zones.

**Figure S3.**
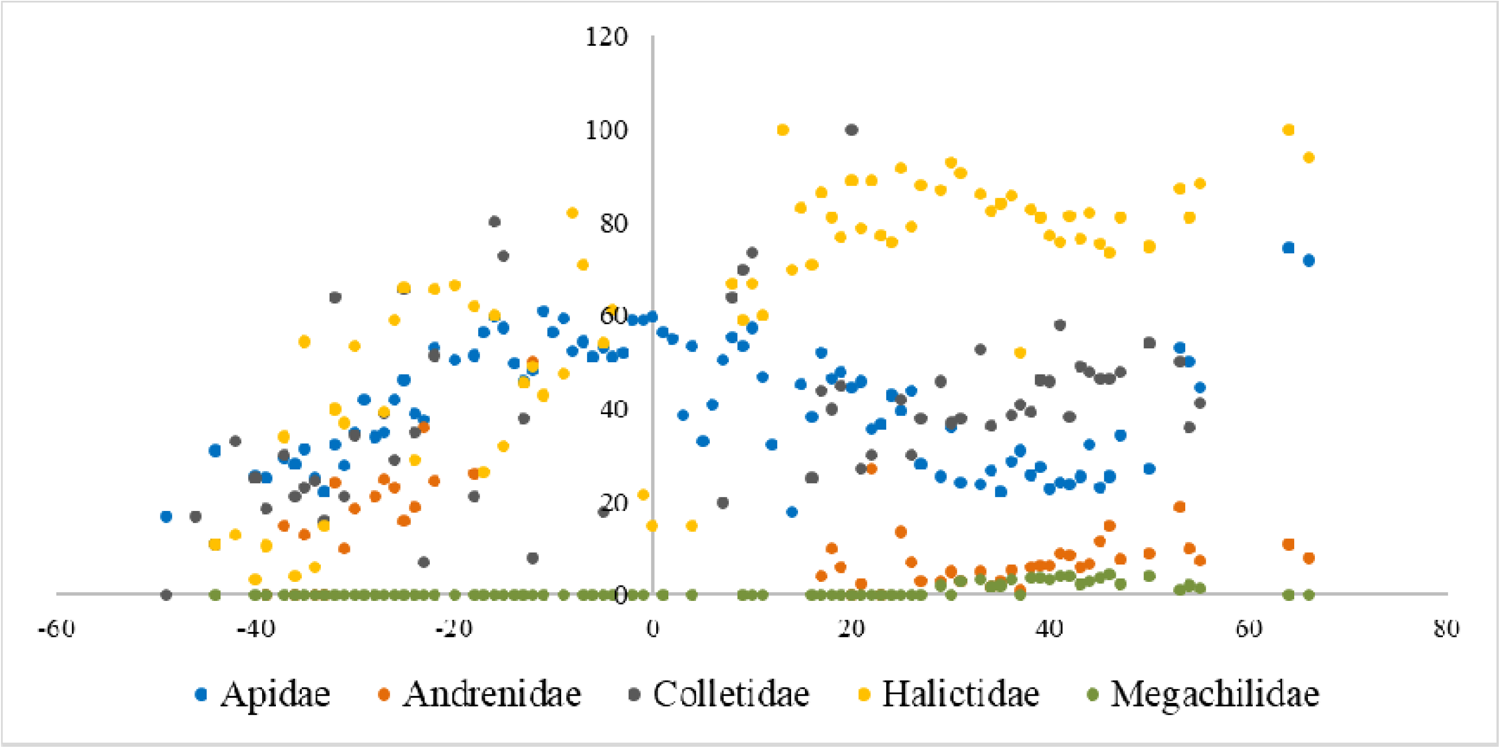
This shows the percentage of all species for all buzzing species (categories 1+2+3) on average with latitude, including all areas with more than 10 species for the Americas. We can see that when a more liberal classification for buzzing is used, the pattern for Halictidae changes, and patterns overall weaken.

###### Europe-Africa

**Figure S4.**
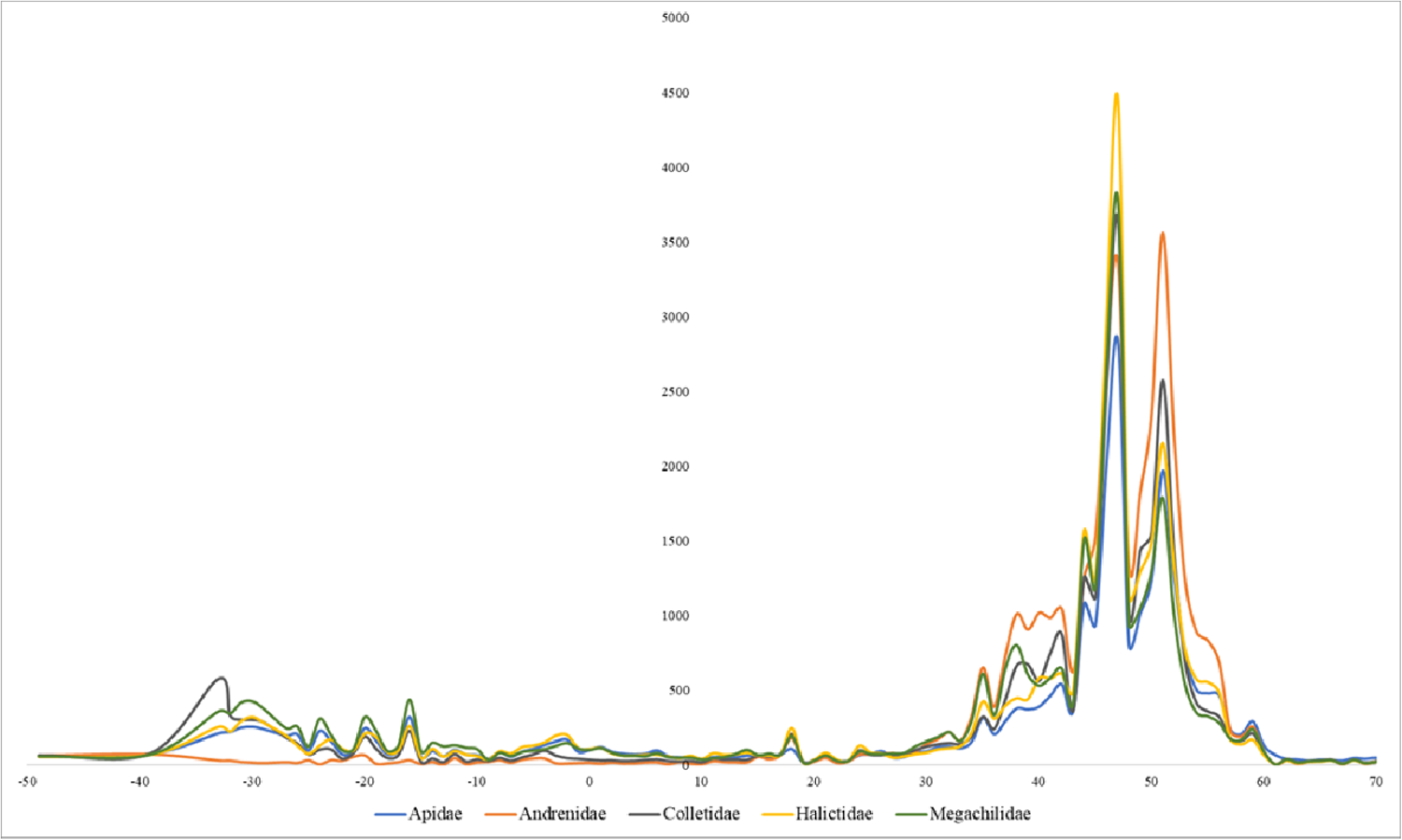
Difference in actual size and distortion based on diversity per unit area between Europe and Africa. Very high diversity in Europe and the Middle East, as well as increasing diversity further south in Africa. However, it should be noted that parts of Africa (particularly sub-Saharan Africa) may have incomplete inventories).

**Figure S5.**
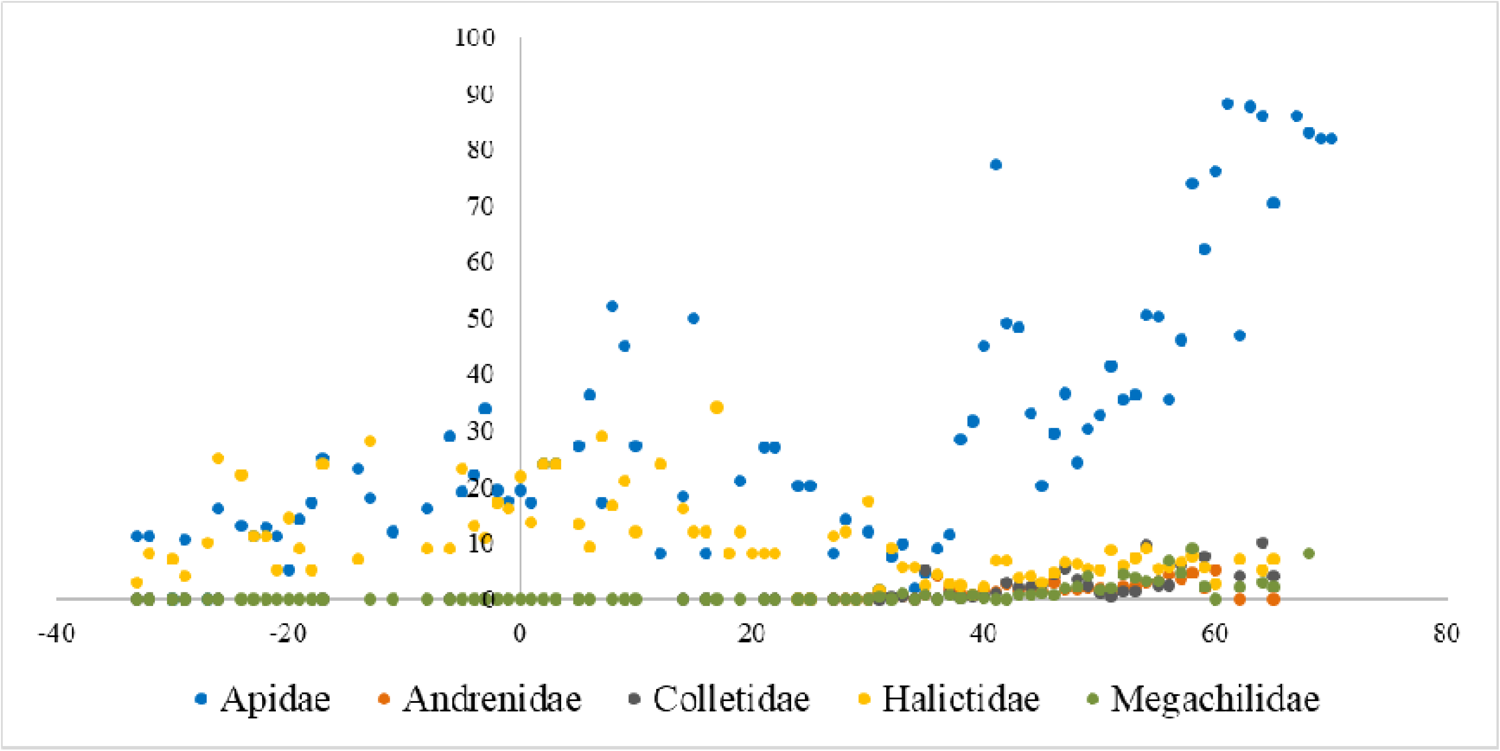
Percentage of species that buzz (categories 1+2) on average with latitude including all areas with more than 10 species between Europe and Africa. Halictidae shows a slight tropical peak in buzzing, whereas Apidae shows a strong temperate peak (*Bombus* alone is provided in Figure S10).

**Figure S6.**
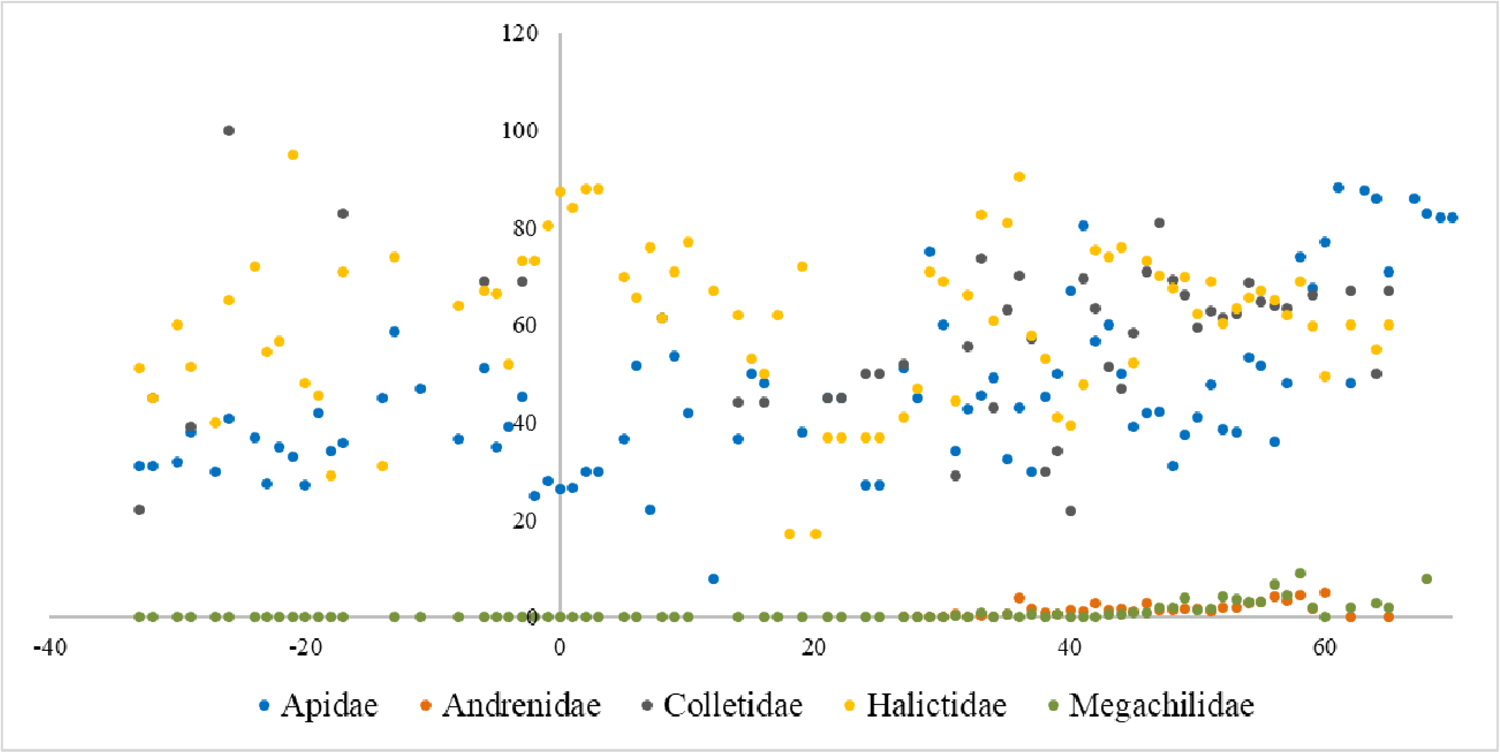
The percentage of species that buzz (categories 1+2+3) on average with latitude including all areas with more than 10 species between Europe and Africa. Adding the third category changes the pattern, but latitudinal trends are overall less clear.

###### Asia-Australia

**Figure S7.**
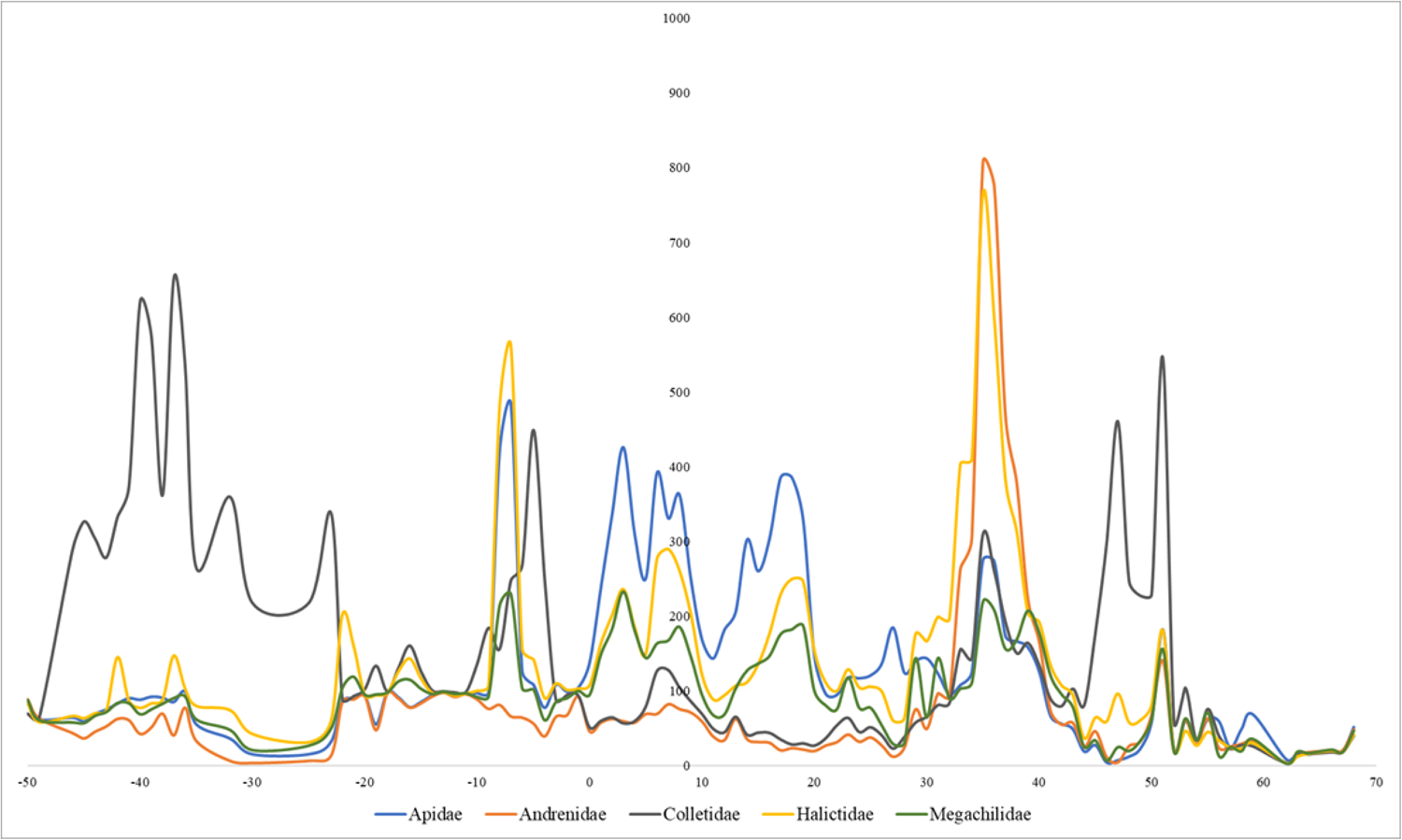
Difference in actual size and distortion based on diversity per unit area for Asia-Australia. Biogeography likely plays a very strong role here, but high richness in Andrenidae and Halictidae at mid-latitudes, Colletidae at high latitudes, and like North America, Apidae at more equatorial latitudes.

**Figure S8.**
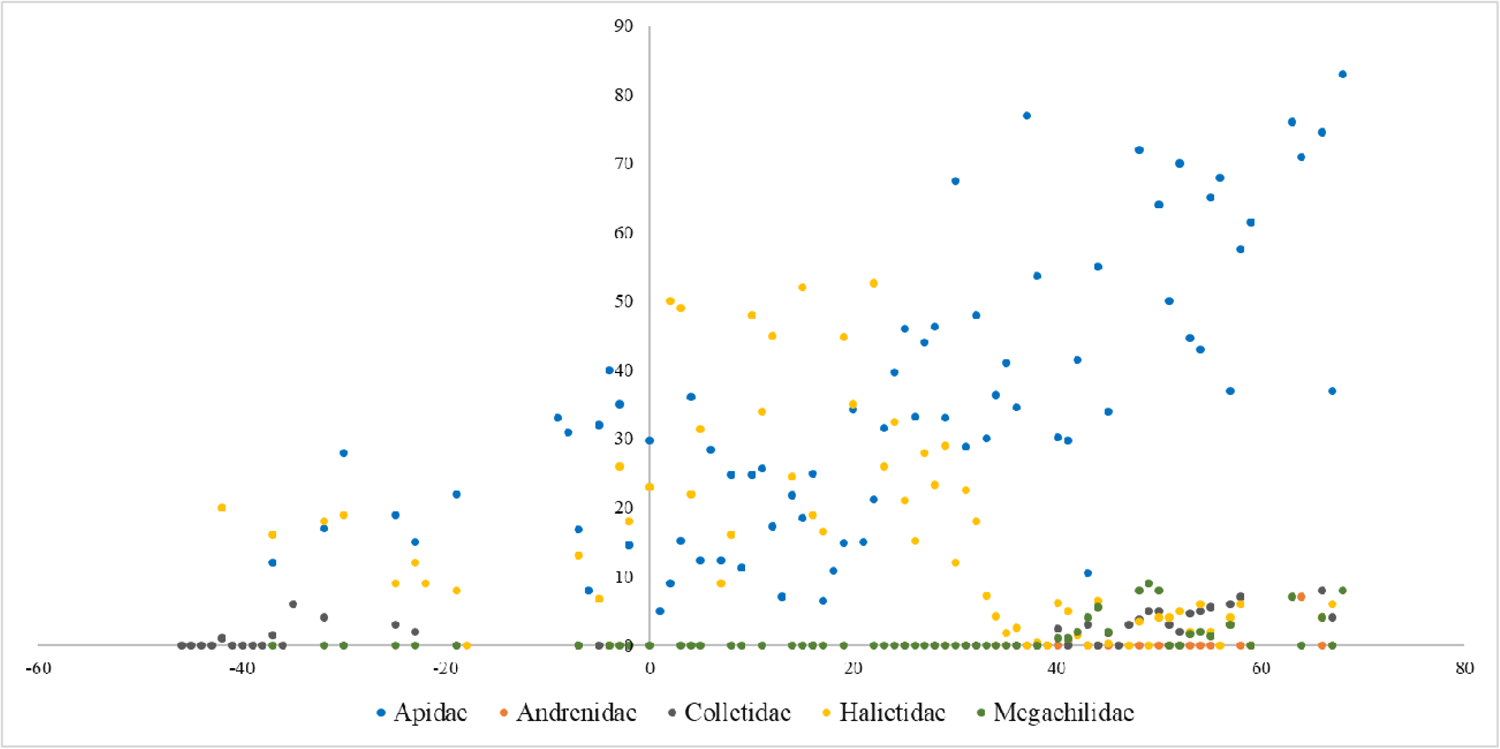
Percentage of species that buzz (categories 1+2) on average with latitude including all areas with more than 10 species for Asia-Australia. Here Halictidae shows a higher percentage in equatorial regions, Apidae at Northern latitudes, and both Megachilidae and Colletidae in temperate zones.

**Figure S9.**
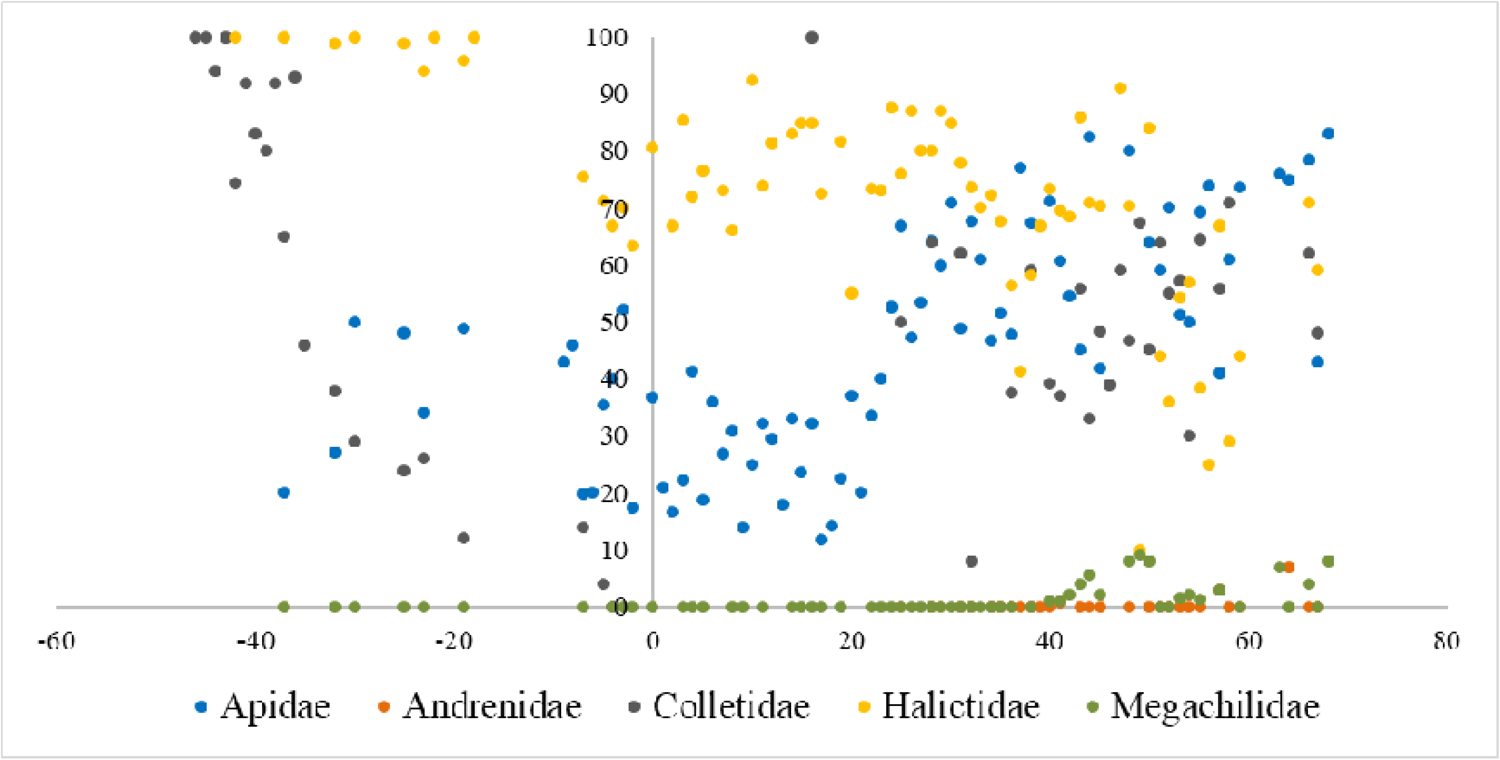
The percentage of all species that buzz (categories 1+2+3) on average with latitude including all areas with more than 10 species for Asia-Australia. Once again the pattern changes when we use the third category to classify buzzing. Here we see much higher percentages of Colletidae showing buzzing at high latitudes in the Northern and Southern Hemispheres, Apidae showing higher buzzing at high latitudes in the Northern Hemisphere (and Andrenidae showing a slight increase in the same region), and Halictidae showing almost the opposite trend to Apidae.

**Figure S10.**
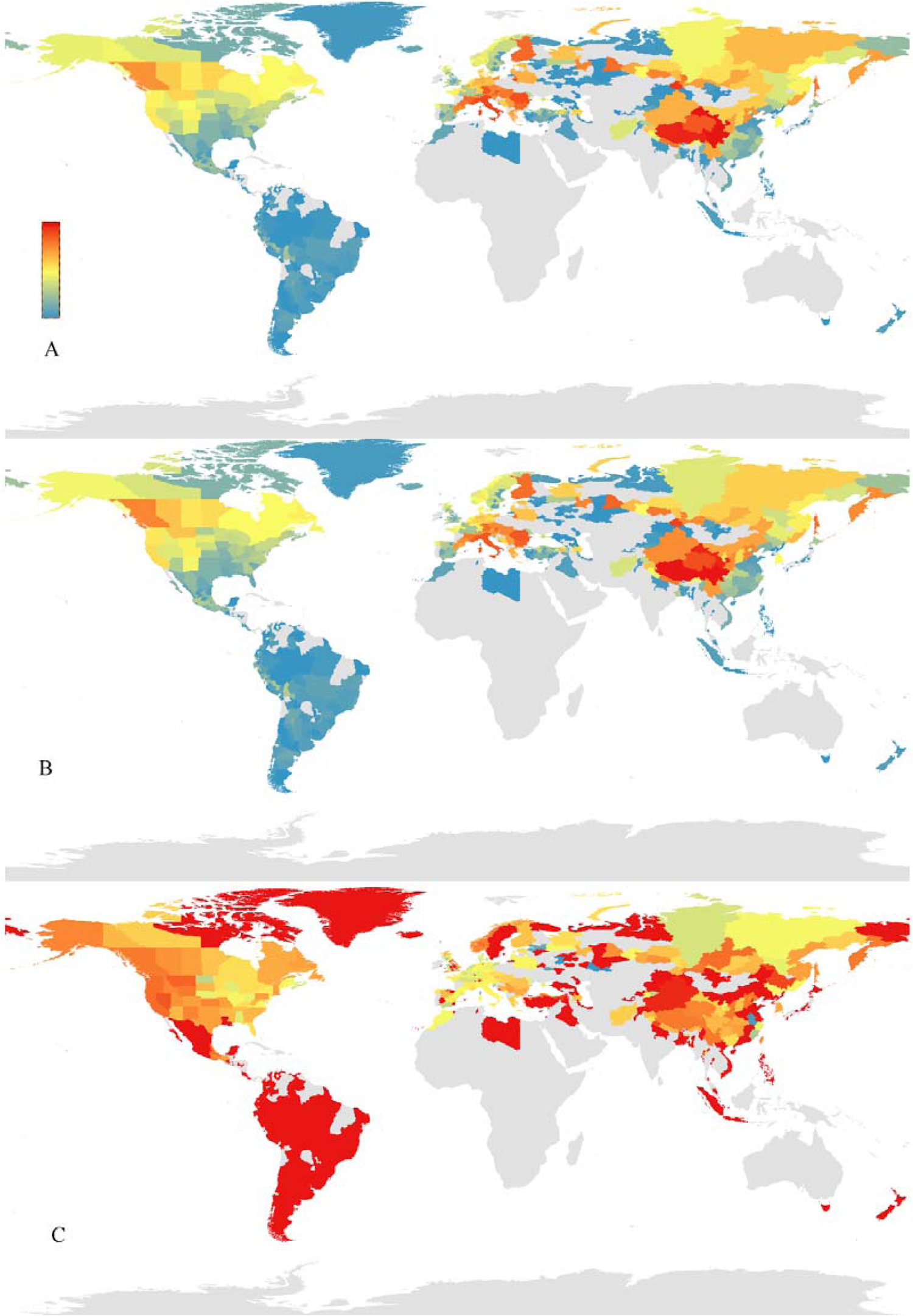
Patterns for bumble bees (*Bombus*), with redder colors indicating higher values, bluer colors indicating lower values, and gray color indicating no species recorded. A. All species richness (max 56); B. Buzz richness, via categories 1+2 (max richness 48); C. Percentage of all species that buzz (categories 1+2+3).

## REFERENCES

Almeida, E.A.B., Bossert, S., Danforth, B.N., Porto, D.S., Freitas, F.V., Davis, C.C., Murray, E.A., Blaimer, B.B., Spasojevic, T., Ströher, P.R., Orr, M.C., Packer, L., Brady, S.G., Kuhlmann, M., Branstetter, M.G., Pie, M.R., 2023. The evolutionary history of bees in time and space. Current Biology 33, 3409–3422.e6. 10.1016/j.cub.2023.07.005

Armbruster, W.S., 2017. The specialization continuum in pollination systems: diversity of concepts and implications for ecology, evolution and conservation. Functional Ecology 31, 88–100. 10.1111/1365-2435.12783

Armbruster, W.S., 2014. Floral specialization and angiosperm diversity: phenotypic divergence, fitness trade-offs and realized pollination accuracy. AoB PLANTS 6. 10.1093/aobpla/plu003

Ascher, J. S., & Pickering, J. (2023). Discover Life bee species guide and world checklist (Hymenoptera: Apoidea: Anthophila).

Bock, C.E., Jones, Z.F., Bock, J.H., 2007. Relationships between species richness, evenness, and abundance in a southwestern savanna. Ecology 88, 1322–1327. 10.1890/06-0654

Brosi, B.J., 2016. Pollinator specialization: from the individual to the community. New Phytologist 210, 1190–1194. 10.1111/nph.13951

Buchmann, S.L., 1985. Bees use vibration to aid pollen collection from non-poricidal flowers. Journal of the Kansas Entomological Society 58, 517–525.

Buchmann, S.L., Hurley, J.P., 1978. A biophysical model for buzz pollination in angiosperms. Journal of Theoretical Biology 72, 639–657. 10.1016/0022-5193(78)90277-1

Buchmann, Stephen L. 1983. Buzz pollination in angiosperms. In: CE Jones, RJ Little, eds. Handbook of experimental pollination biology. New York, NY, USA: Scientific and Academic Editions, 73–113.

Cameron, S.A., Sadd, B.M., 2020. Global trends in bumble bee health. Annu. Rev. Entomol. 65, 209–232. 10.1146/annurev-ento-011118-111847

Cardinal, S., Buchmann, S.L., Russell, A.L., 2018. The evolution of floral sonication, a pollen foraging behavior used by bees (Anthophila). Evolution 72, 590–600. 10.1111/evo.13446

Cariveau, D.P., Winfree, R., 2015. Causes of variation in wild bee responses to anthropogenic drivers. Current Opinion in Insect Science 10, 104–109. 10.1016/j.cois.2015.05.004

Chesshire, P.R., Fischer, E.E., Dowdy, N.J., Griswold, T.L., Hughes, A.C., Orr, M.C., Ascher, J.S., Guzman, L.M., Hung, K.J., Cobb, N.S., McCabe, L.M., 2023. Completeness analysis for over 3000 United States bee species identifies persistent data gap. Ecography 2023, e06584. 10.1111/ecog.06584

Cooley, H., Vallejo-Marín, M., 2021. Buzz-pollinated crops: a global review and meta-analysis of the effects of supplemental bee pollination in tomato. Journal of Economic Entomology 114, 505–519. 10.1093/jee/toab009

Corbet, S.A., Beament, J., Eisikowitch, D., 1982. Are electrostatic forces involved in pollen transfer? Plant Cell & Environment 5, 125–129. 10.1111/1365-3040.ep11571488

Corbet, S.A., Huang, S.-Q., 2014. Buzz pollination in eight bumblebee-pollinated *Pedicularis* species: does it involve vibration-induced triboelectric charging of pollen grains? Annals of Botany 114, 1665–1674. 10.1093/aob/mcu195

Denelle, P., Weigelt, P., & Kreft, H. 2023. GIFT—An R package to access the global inventory of floras and traits. Methods in Ecology and Evolution 14 (11), 2738.

De Luca, P.A., Vallejo-Marín, M., 2013. What’s the ‘buzz’ about? The ecology and evolutionary significance of buzz-pollination. Current Opinion in Plant Biology 16, 429–435. 10.1016/j.pbi.2013.05.002

Delgado, T., Leal, L.C., El Ottra, J.H.L., Brito, V.L.G., Nogueira, A., 2023. Flower size affects bee species visitation pattern on flowers with poricidal anthers across pollination studies. Flora 299, 152198. 10.1016/j.flora.2022.152198

Dorey, James B. 2023. A globally synthesised and flagged bee occurrence dataset and cleaning workflow. Scientific Data 10, 747. 10.1038/s41597-023-02626-w

Galen, C., 1999. Why Do Flowers Vary? BioScience 49, 631–640. 10.2307/1313439

Galen, C., Zimmer, K.A., Newport, M.E., 1987. Pollination in floral scent morphs of *Polemonium viscosum*lJ: a mechanism for disruptive selection on flower size. Evolution 41, 599–606. 10.1111/j.1558-5646.1987.tb05830.x

González-Vanegas, P.A., Rös, M., García-Franco, J.G., Aguirre-Jaimes, A., 2021. Buzz-pollination in a tropical montane cloud forest: compositional similarity and plant-pollinator interactions. Neotrop Entomol 50, 524–536. 10.1007/s13744-021-00867-1

Houston, T.F., Ladd, P.G., 2002. Buzz pollination in the Epacridaceae. Aust. J. Bot. 50, 83. 10.1071/BT01020

Irwin, R.E., Cook, D., Richardson, L.L., Manson, J.S., Gardner, D.R., 2014. Secondary Compounds in Floral Rewards of Toxic Rangeland Plants: Impacts on Pollinators. J. Agric. Food Chem. 62, 7335–7344. 10.1021/jf500521w

Klomberg, Y., Dywou Kouede, R., Bartoš, M., Mertens, J.E.J., Tropek, R., Fokam, E.B., Janeček, Š., 2019. The role of ultraviolet reflectance and pattern in the pollination system of *Hypoxis camerooniana* (Hypoxidaceae). AoB PLANTS 11, plz057. 10.1093/aobpla/plz057

Knight, J., 2022. Scientists’ warning of the impacts of climate change on mountains. PeerJ 10, e14253. 10.7717/peerj.14253

Koski, M.H., Ashman, T., 2014. Dissecting pollinator responses to a ubiquitous ultraviolet floral pattern in the wild. Functional Ecology 28, 868–877. 10.1111/1365-2435.12242

Koski, M.H., MacQueen, D., Ashman, T.-L., 2020. Floral pigmentation has responded rapidly to global change in ozone and temperature. Current Biology 30, 4425–4431.e3. 10.1016/j.cub.2020.08.077

Leclercq, N., Marshall, L., Caruso, G., Schiel, K., Weekers, T., Carvalheiro, L.G., Dathe, H.H., Kuhlmann, M., Michez, D., Potts, S.G., Rasmont, P., Roberts, S.P.M., Smagghe, G., Vandamme, P., Vereecken, N.J., 2023. European bee diversity: Taxonomic and phylogenetic patterns. Journal of Biogeography 50, 1244–1256. 10.1111/jbi.14614

LeCroy, K.A., Savoy-Burke, G., Carr, D.E., Delaney, D.A., Roulston, T.H., 2020. Decline of six native mason bee species following the arrival of an exotic congener. Sci Rep 10, 18745. 10.1038/s41598-020-75566-9

Liu, Y., Xu, X., Dimitrov, D., Pellissier, L., Borregaard, M.K., Shrestha, N., Su, X., Luo, A., Zimmermann, N.E., Rahbek, C., Wang, Z., 2023. An updated floristic map of the world. Nat Commun 14, 2990. 10.1038/s41467-023-38375-y

Lozada-Gobilard, S., Motter, A., Sapir, Y., 2023. AmongLyears rain variation is associated with flower size, but not with signal patch size in *Iris petrana*. Ecology 104, e3839. 10.1002/ecy.3839

Mesquita-Neto, J.N., Blüthgen, N., Schlindwein, C., 2018. Flowers with poricidal anthers and their complex interaction networks—Disentangling legitimate pollinators and illegitimate visitors. Functional Ecology 32, 2321–2332. 10.1111/1365-2435.13204

Olofsson, J., Shams, H., 2007. Determinants of plant species richness in an alpine meadow. Journal of Ecology 95, 916–925. 10.1111/j.1365-2745.2007.01284.x

Orr, M.C., Hughes, A.C., Chesters, D., Pickering, J., Zhu, C.-D., Ascher, J.S., 2021. Global Patterns and Drivers of Bee Distribution. Current Biology 31, 451–458.e4. 10.1016/j.cub.2020.10.053

Pacheco Filho, A.J.D.S., Verola, C.F., Lima Verde, L.W., Freitas, B.M., 2015. Bee-flower association in the Neotropics: implications to bee conservation and plant pollination. Apidologie 46, 530–541. 10.1007/s13592-014-0344-8

Page, M.L., Williams, N.M., 2023. Evidence of exploitative competition between honey bees and native bees in two California landscapes. Journal of Animal Ecology 92, 1802–1814. 10.1111/1365-2656.13973

Palmer-Young, E.C., Farrell, I.W., Adler, L.S., Milano, N.J., Egan, P.A., Junker, R.R., Irwin, R.E., Stevenson, P.C., 2019. Chemistry of floral rewards: intraL and interspecific variability of nectar and pollen secondary metabolites across taxa. Ecological Monographs 89, e01335. 10.1002/ecm.1335

Pemberton, R.W., 2023. Plant Resource Use and Pattern of Usage by the Naturalized Orchid Bee (*Euglossa dilemma*: Hymenoptera: Apidae) in Florida. Insects 14, 909. 10.3390/insects14120909

Potts, S. G., Biesmeijer, J. C., Kremen, C., Neumann, P., Schweiger, O., & Kunin, W. E. 2010. Global pollinator declines: trends, impacts and drivers. Trends in ecology & evolution, 25(6), 345–353.

Roulston, T.H., Cane, J.H. 2000. Pollen nutritional content and digestibility for animals. Plant Systematics and Evolution 222, 187–209. 10.1007/BF00984102

Russell, A.L., Buchmann, S.L., Papaj, D.R., 2017. How a generalist bee achieves high efficiency of pollen collection on diverse floral resources. Behavioral Ecology 28, 991–1003. 10.1093/beheco/arx058

Russell, A.L., Papaj, D.R., 2016. Artificial pollen dispensing flowers and feeders for bee behaviour experiments. J Poll Ecol 18, 13–22. 10.26786/1920-7603(2016)14

Russell, A.L., Buchmann, S.L., Zenil-Ferguson, R., Jolles, D., Kriebel, R, Vallejo-Marín, M. 2024. Widespread evolution of poricidal flowers: A striking example of morphological convergence across flowering plants. BioRxiv. 10.1101/2024.02.28.582636

Schlindwein, C. 2004. Are oligolectic bees always the most effective pollinators? In: Freitas, B.M., Pereira, J.O.P. (eds.) Solitary Bees Conservation, Rearing and Management for Pollination, pp. 231–240. Federal University of Ceará, Fortaleza

Song, B., Sun, L., Barrett, S.C.H., Moles, A.T., Luo, Y., Armbruster, W.S., Gao, Y., Zhang, S., Zhang, Z., Sun, H., 2022. Global analysis of floral longevity reveals latitudinal gradients and biotic and abiotic correlates. New Phytologist 235, 2054–2065. 10.1111/nph.18271

Sutter, L., Jeanneret, P., Bartual, A.M., Bocci, G., Albrecht, M., 2017. Enhancing plant diversity in agricultural landscapes promotes both rare bees and dominant cropLpollinating bees through complementary increase in key floral resources. Journal of Applied Ecology 54, 1856–1864. 10.1111/1365-2664.12907

Vallejo-Marín, M., 2022. How and why do bees buzz? Implications for buzz pollination. Journal of Experimental Botany 73, 1080–1092. 10.1093/jxb/erab428

Vallejo-Marin, M., Russell, A.L., 2023. Harvesting pollen with vibrations: towards an integrative understanding of the proximate and ultimate reasons for buzz pollination. Annals of Botany mcad189. 10.1093/aob/mcad189

Van Der Kooi, C.J., Vallejo-Marín, M., Leonhardt, S.D., 2021. Mutualisms and (A)symmetry in Plant–Pollinator Interactions. Current Biology 31, R91–R99. 10.1016/j.cub.2020.11.020

Vanderplanck, M., Decleves, S., Roger, N., Decroo, C., Caulier, G., Glauser, G., Gerbaux, P., Lognay, G., Richel, A., Escaravage, N., Michez, D., 2018. Is nonLhost pollen suitable for generalist bumblebees? Insect Science 25, 259–272. 10.1111/1744-7917.12410

Villalobos, S., Sevenello-Montagner, J.M., Vamosi, J.C., 2019. Specialization in plant–pollinator networks: insights from local-scale interactions in Glenbow Ranch Provincial Park in Alberta, Canada. BMC Ecol 19, 34. 10.1186/s12898-019-0250-z

Vit, P., Pedro, S.R.M., Roubik, D.W. (Eds.), 2018. Pot-Pollen in Stingless Bee Melittology. Springer International Publishing, Cham. 10.1007/978-3-319-61839-5

Winfree, R., 2010. The conservation and restoration of wild bees. Annals of the New York Academy of Sciences 1195, 169–197. 10.1111/j.1749-6632.2010.05449.x

Xie, L., Guo, H., Ma, C., 2016. Alterations in flowering strategies and sexual allocation of *Caragana stenophylla* along a climatic aridity gradient. Sci Rep 6, 33602. 10.1038/srep33602

